# Mechanical Strategies Supporting Growth and Size Diversity in Filamentous Fungi

**DOI:** 10.1101/2024.04.17.589919

**Authors:** Louis Chevalier, Flora Klingelschmitt, Ludovic Mousseron, Nicolas Minc

## Abstract

The stereotypical tip growth of filamentous fungi supports their lifestyles and functions. It relies on the polarized remodeling and expansion of a protective elastic cell wall (CW) driven by large cytoplasmic turgor pressure. Remarkably, hyphal filament diameters, and cell elongation rates can vary extensively among different fungi. To date, however, how fungal cell mechanics may be adapted to support these morphological diversities while ensuring surface integrity remains unknown. Here, we combined super-resolution imaging and deflation assays, to measure local CW thickness, elasticity and turgor in a set of fungal species spread on the evolutionary tree that span a large range in cell size and growth speeds. While CW elasticity exhibited dispersed values presumably reflecting differences in CW composition, both thickness and turgor scaled in dose-dependence with cell diameter and growth speeds. Notably, larger cells exhibited thinner lateral CWs, and faster cells thinner apical CWs. Counterintuitively, turgor pressure was also inversely scaled with cell diameter and tip growth speed, challenging the idea that turgor is the primary factor dictating tip elongation rates. We propose that fast growing cells with rapid CW turnover have evolved strategies based on a less turgid cytoplasm and thin walls to safeguard wall integrity and survival.

## INTRODUCTION

Survival is a fundamental constraint that shapes the evolution of living organisms. The survival, life styles and function of many organisms, including bacteria, fungi and plants, is associated to the synthesis of a rigid cell wall (CW) that encases the plasma membrane to mechanically protect these cells. CWs are composed of diverse polysaccharides and proteins, that largely vary among species and phyla. CWs are elastic and as such maintain cell shapes, but they also remodel locally to incorporate plastic deformation, a process key for cell growth (1, 2). Walled cells also feature a large internal cytoplasmic turgor pressure which is osmotically generated, that reach values on the order of several atmospheres (3–5). Turgor inflates the CW and put it under tension, and is as such thought to be a key driving element to permit growth against fundamental forces including gravity and resistive forces from multiple substrates such as soils or host tissues (2, 6, 7). In general, however, how walled cell mechanics may be fit to the function, morphology or environment of different organisms remain poorly studied.

Filamentous fungi are ancestral walled eukaryotes, which span ∼600 Myears of evolution (8). They occupy most environments on earth, and represent an important class of pathogens for animals and plants. A wide spread strategy adopted by fungi to colonize space, reproduce or infect their hosts is tip growth (9). Tip growth implicates a highly polarized synthesis and remodeling of the CW at cell tips, in part regulated by the cytoskeleton and small GTPases like Cdc42 and Rac that restrict the secretion of synthesis and remodeling CW enzymes to cell tips (1, 9–11). Interestingly, in spite of conserved mechanisms and effectors, tip growth and hyphal shapes feature a remarkable variability across species, with some fungi like *Fusarium oxyparum*, that grow thin hyphae of less than 1 µm in diameter (12), up to very large cells found in *Neurospora crassa* or *Sclerotinia sclerotiorum* that may reach diameters as large as 10-20 µm (13). Tip growth speeds also span a remarkable large range of values, from relatively slow elongating yeast cells that grow at ∼1µm/hr up to some of the fastest fungi that elongate as fast as 500 µm/hr (14, 15).

Variations in growth speeds and hyphal shapes may derive from the modulation of rates of CW synthesis and turnover, and on the size of polarity/secretion zones, respectively (16–19). However, deformation of freshly assembled wall portions is also thought to require tensional stresses derived from turgor, as reducing turgor in most fungal cells halts growth in a near-instantaneous manner (3, 20). Importantly, such tensional stresses also entail large risks of CW failure, with CW elastic strains at cell tips estimated to be around 20-30% and failure strains, at which the CW breaks open, measured to be around ∼40-50% in some fungal species (21, 22).

Accordingly, tip bursting is a common phenotype of fungal hyphae treated with hypoosmotic medium, to increase turgor pressure, or of mutant cells defective in CW synthesis (23, 24). To date, which mechanical strategies fungal cells may have employed to cope with the risks associated with rapid cell growth have not been tackled in a systematic manner. In part progress in this fundamental question has been limited by the low-throughput associated with typical methods like AFM, commonly used to measure CW mechanics and turgor, and that of electron microscopy to measure CW thickness.

Here, we adapted a sub-resolution method to image CW thickness (16, 24) in live cells of 7 fungal species selected among distant branches on the fungal evolutionary tree, based on their differences in hyphal cell diameters and elongation speeds. We coupled thickness measurements to strain-stress assays based on CW elastic relaxation achieved by ablating CWs with a laser and/or the application of ranges of osmotic shocks (21, 25). This allowed to measure with medium throughput basal values of turgor pressure, CW thickness and elastic moduli. Our results reveal relatively dispersed values of elastic moduli both at cell tips and sides, plausibly reflecting differences in CW compositions, across fungi. In contrast, both CW thickness and turgor appeared to inversely scale with cell size and tip growth. These findings suggest a counterintuitive scaling with rapid growing and large cells featuring less tensed cell walls. We propose that this strategy may limit risks of failure when CWs remodel rapidly.

## RESULTS AND DISCUSSIONS

### Mapping the mechanical properties in different fungal species

To analyze how CW mechanics and turgor may vary among fungal species that feature cells with different sizes and elongation speeds, we selected 7 representative species across the fungal kingdom, namely *Neurospora crassa, Botrytis cinerea, Aspergillus nidulans, Penicilium janczewskii, Candida albicans, Schizosaccharomyces pombe and Mucor circinelloides* (Fig 1A and Table S1). Measurements for *S. Pombe* and *A. nidulans* were taken from our own previous studies employing the exact same strategy as described here (16, 21). Among these species, 6 are septate fungi that belong to different clades of ascomycetes, and one, *Mucor circinelloides* is a more ancient non-septate basidomycetes fungus. Also, several of these selected fungi are known opportunistic human pathogens (*Mucor circinelloides and Candida albicans*), and one is a plant pathogen (*Botrytis cinerea*). Species were selected based on morphological features, especially hyphal diameter and tip elongation speeds from previous reports, and also as relatively simple systems to grow in laboratory conditions (13, 14) (Fig 1B). Importantly, hyphal cells were all imaged using similar microscopy set ups but media and temperature were adapted to the different species (Table S1).

**Figure 1:**
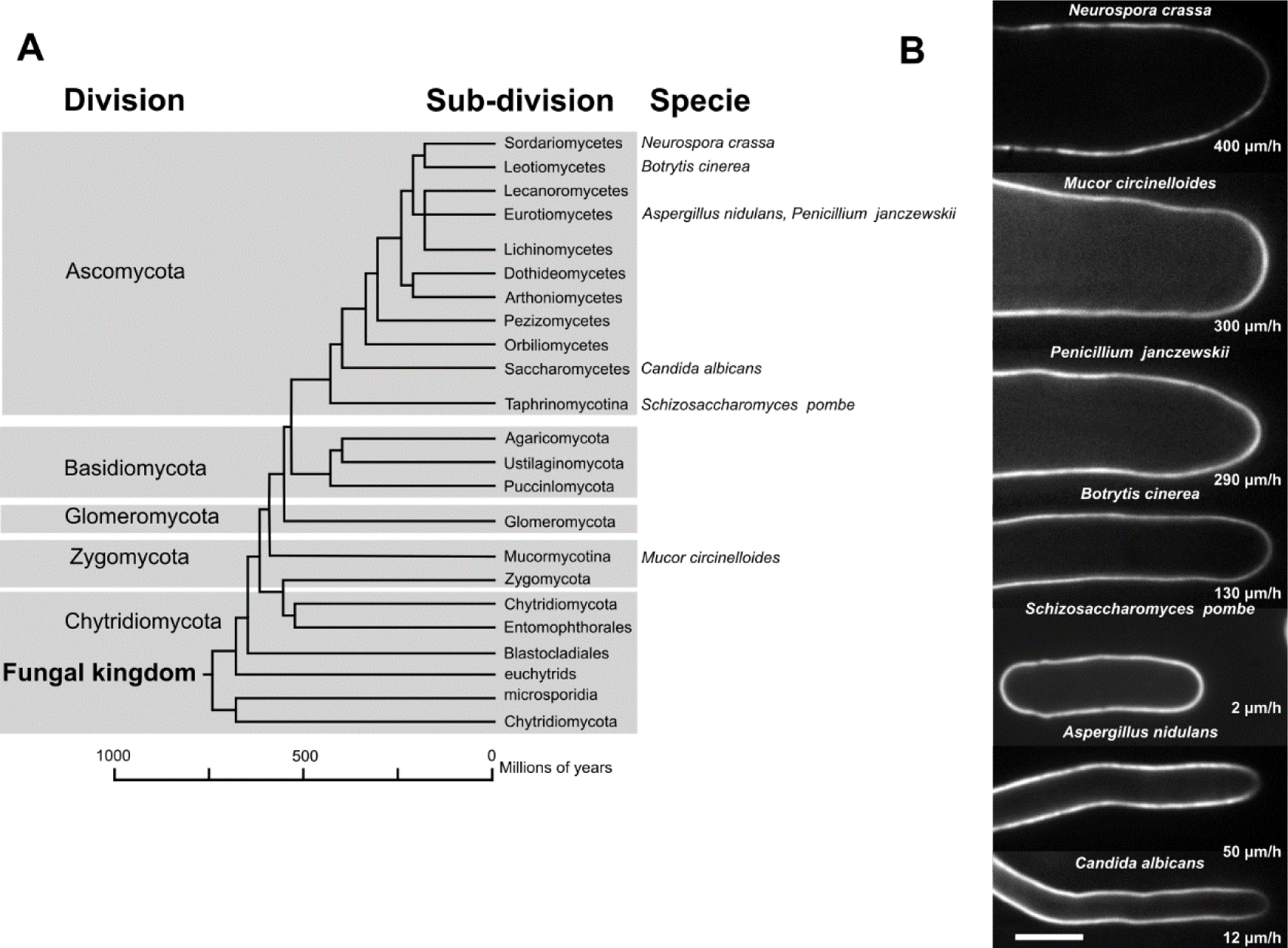
Diversity of size and speed across the fungal kingdom. (A) Phylogenetic tree showing fungal division, sub-division, and species used here. Adapted from (42, 43). (B) Fluorescent confocal images of lectin signal to label the CW surface in the different used species, and their mean growth speed. Scale bar =5µm.

In order to measure CW thickness, h, around live hyphae, we adapted a previous sub-resolution method developed in *S. Pombe* and *A. nidulans*, based on the sub-pixel segmentation of the inner and outer faces of CWs, labelled with fluorophores of different colors (16, 21). We labeled the outer face of CWs with either WGA, ConA or GS-IB4 lectins (depending on species) bound to far-red emitting fluorophores (Fig S1 and Table S2). Membranes were labeled with integral or membrane-bound proteins in tractable species (*Schizosaccharomyces pombe*, *Aspergillus nidulans, Candida albicans* and *Neurospora crassa*) (16, 24, 26, 27), and by using the FM-4 64 dye in other species (Fig S1 and Table S2) (13). However, as this dye is rapidly endocytosed, cells were placed in open liquid chambers, labeled with lectins, and FM-4 64 was added close to the field of view before immediate imaging. After imaging membranes and lectins, cells were rapidly pierced using a UV laser focused on the cell surface. This caused cells to deflate and CWs to relax, from which we measured CW elastic strains (21, 25). Both local thickness and elastic strains allowed to estimate local values of CW elastic moduli divided by pressure, Y/P, on cell sides or at cell tips (21). Finally, CW elastic strains measured from laser ablation were compared with elastic strains at different concentrations of sorbitol added to the medium, and media osmolarity was measured with an osmometer, to estimate turgor values, P (Fig S2) (25). Therefore, these methodologies allowed to compute key local and global mechanical parameters at the single cell level in multiple filamentous fungi (Fig 2A).

**Figure 2:**
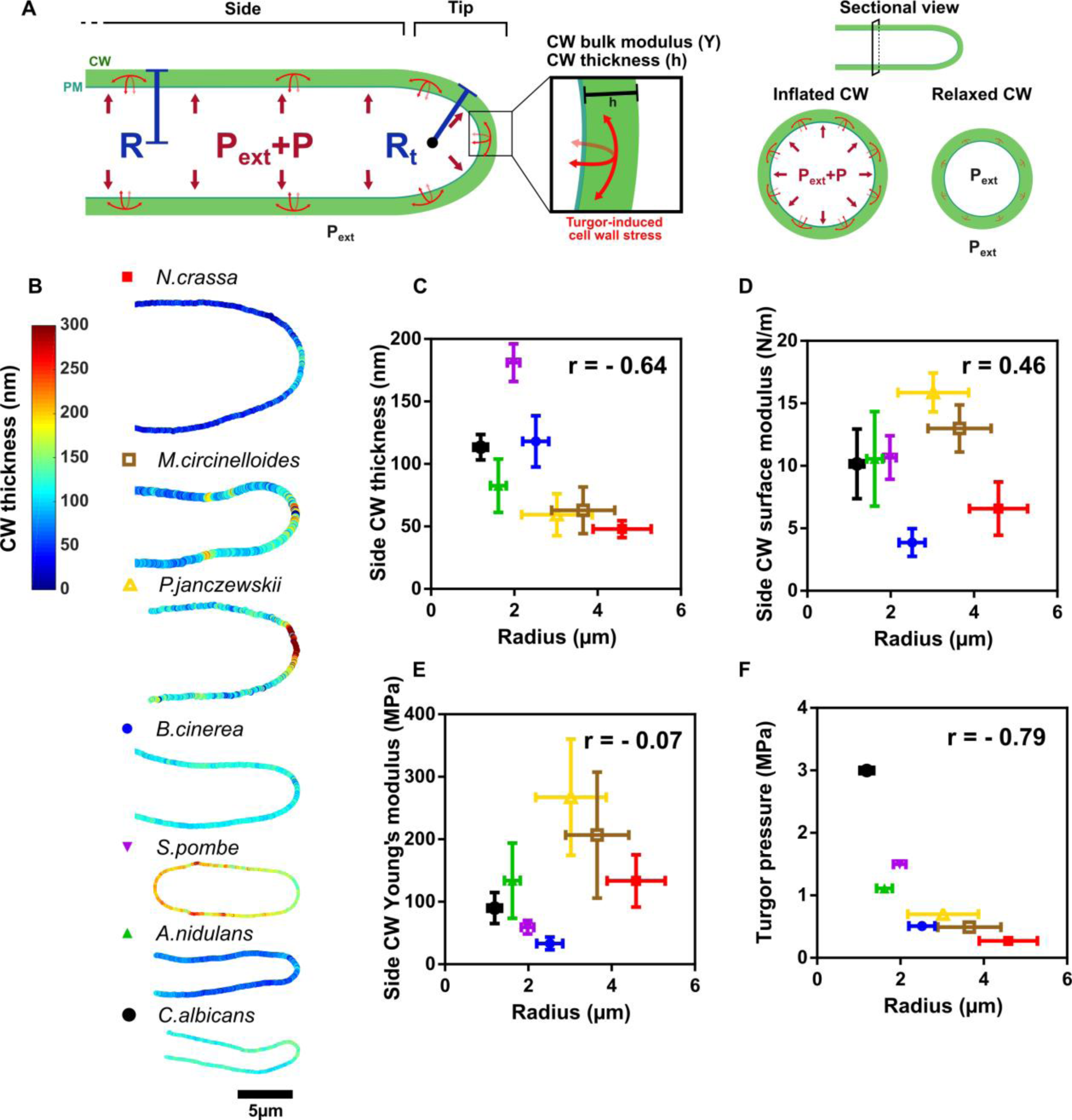
Cell mechanics and cell size in filamentous fungi. (A) Scheme of hyphal mechanics, representing the radius (R), the tip curvature radius (Rt), the turgor pressure (P), the cell wall thickness (h), and the CW Young’s modulus (Y). Internal turgor pressure stretches elastically the CW. (B) Color maps of CW thickness for the different fungal species considered in this study. (C-F) Mechanical parameters of lateral CWs plotted against the cell radius. (C) Side cell wall thickness (n>16 cells for each species), (D) side CW Young’s modulus (n>16 for each species), (E) side CW surface modulus (n>11 for each species), and (F) turgor pressure (n>15 for each species). Error bars correspond to +/- S.D. Indicated r values on the graphs are Pearson correlation coefficients. Scale bar =5µm.

### Scaling of cell mechanical parameters with hyphal cell diameter

We first explored putative correlations between cell radius, R, CW mechanical properties and turgor. In the fungi considered here, values of thicknesses of lateral CWs ranged from ∼150-200nm down to 30-50nm, highlighting large variations among different species (Fig 2B-2C). Interestingly, lateral CW thickness values were generally smaller in wider cells for all the species considered, although we noted that *S. Pombe* was above the general trend of other fungi. Importantly, the values measured agreed with thickness measurements from electron microscopy images found in the literature, when available (Fig 2B-2C) (24, 28–31). In contrast, elastic Young’s moduli that reflect bulk material properties of CWs and their polysaccharide composition and arrangements were more dispersed (Fig 2E). We interpret these variations as a plausible result of large variations in CW composition among fungi (32, 33). CW surface moduli computed as the product of h and Y, which reflect an apparent stiffness for the CW, did not appear to increase with cell diameters and was also dispersed (Fig 2D). This suggests that larger lateral CWs are not necessarily stiffer among different fungal species. Strikingly, turgor values also showed clear dose-dependence changes with cell diameters, but surprisingly, wider cells appeared to feature a less turgid cytoplasm (Fig 2F). We conclude that fungal cells exhibit marked scaled variations in the mechanical properties of their lateral CWs as well as turgor values with their hyphal geometry, with larger cells having thinner walls and being less turgid.

To understand how these values may functionally affect the mechanical state of lateral CWs, we computed CW tension (PR), stress (PR/h), elastic strain (PR/hY), and a bending energy rescaled to that of pressure (PR^3^/Yh^3^) (Fig S3) (21). This analysis showed that CW tension was generally lower in larger cells, but did not reveal any simple scaling relationships for CW stress (Fig S3A and S3B). Rescaled bending energy values also increased progressively with cell diameter, suggesting that larger lateral CWs found in different species may be easier to bend (Fig S3D). Finally, with the exception of *Candida albicans,* which was above the general trend, the elastic strain of lateral CWs was mostly flat, at values around ∼20%, raising the possibility that limitations in CW deformation may constrain cell diameter definition to ensure wall integrity. We conclude that large fungal cells feature less pressurized cytoplasm and thinner lateral CWs, which contribute to limit elastic strains in their larger lateral CWs.

### Scaling of cell mechanical parameters with hyphal cell growth

Next, we focused on CW mechanical properties at cell tips, to understand how they may be modulated as a function of hyphal cell elongation speeds. As previously reported, larger cells generally grew faster (14); presumably reflecting a higher CW remodeling activity at cell tips in larger cells (15) (Fig 3A). Interestingly, we found that values of CW thickness at cell tips were inversely correlated with growth speeds, suggesting that rapid growing cells feature thinner tip CWs (Fig 3B). In addition, the ratio of CW thickness between cell tips and cell sides also increased with tip growth speeds, with rapid species like *P. janczewskii, M. circinelloides* and *N. crassa* that featured a thicker CW at cell tips as compared to cell sides, and relatively slower ones like *A. nidulans* and *S. pombe*, with the opposite CW thickness polarity (Fig 4A). Both CW Young’s and surface elastic moduli at cell tips, exhibited dispersed values among different species as for lateral CWs (Fig 3C-3D). Yet, the ratio of both bulk and surface moduli between tips and lateral CWs was systematically below 1, in agreement with the notion that CW at cell tips are softer than on cell sides for an anisotropic cell elongation (Fig 4B-C) (7). In addition, this ratio was smaller in faster growing cells, indicating that the polarization of CW stiffness may become more pronounced as cells grow faster (Fig 4B-C). Finally, one of the most unique and surprising features was to find that faster growing cells have lower turgor pressure values than slow growing ones (Fig 3E). For instance, fission yeast turgor was measured to be around 1.5MPa with cells elongating at ∼2 µm/h and turgor in *N. crassa* was only ∼0.2MPa with cells growing at ∼400 µm/h. Together, these findings suggest a counterintuitive scaling in which faster growing cells feature thinner walls at their cell tips and a less turgid cytoplasm.

**Figure 3:**
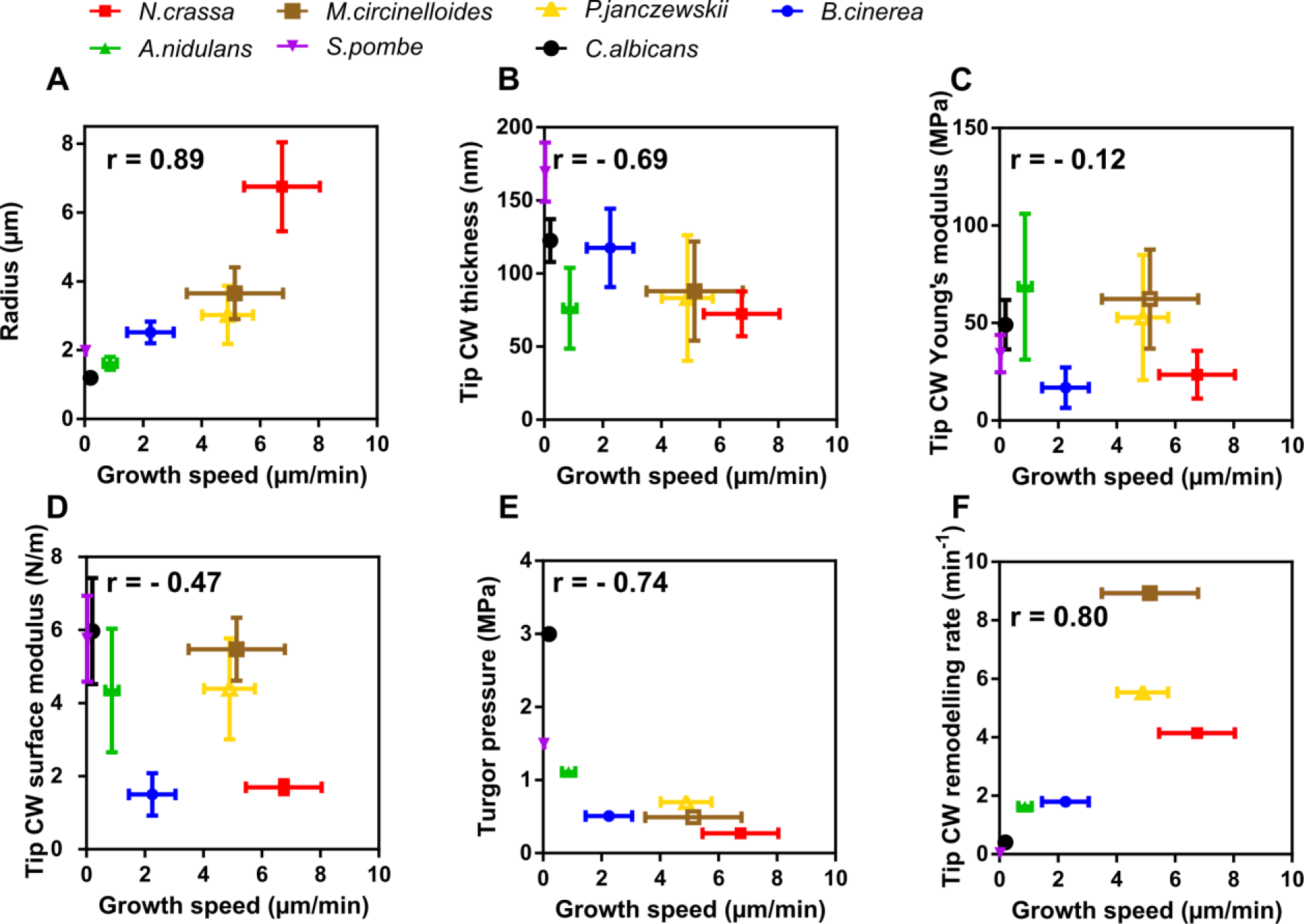
Apical mechanical parameters dependence on tip growth speed. Mechanical and geometrical parameters plotted against average growth speed for different fungal species. (A) Radius (n>23 cells for each specie), (B) tip CW thickness (n>11 for each specie), (C) tip CW Young’s modulus (n>11 for each specie), (D) tip CW surface modulus (n>7 for each specie), (E) pressure (n>15 for each specie), and (F) Effective CW remodeling rate. The effective remodeling rate is calculated as the growth speed divided by PR²/Yh. Error bars correspond to +/- S.D. Indicated r values on the graphs are Pearson correlation coefficients.

**Figure 4:**
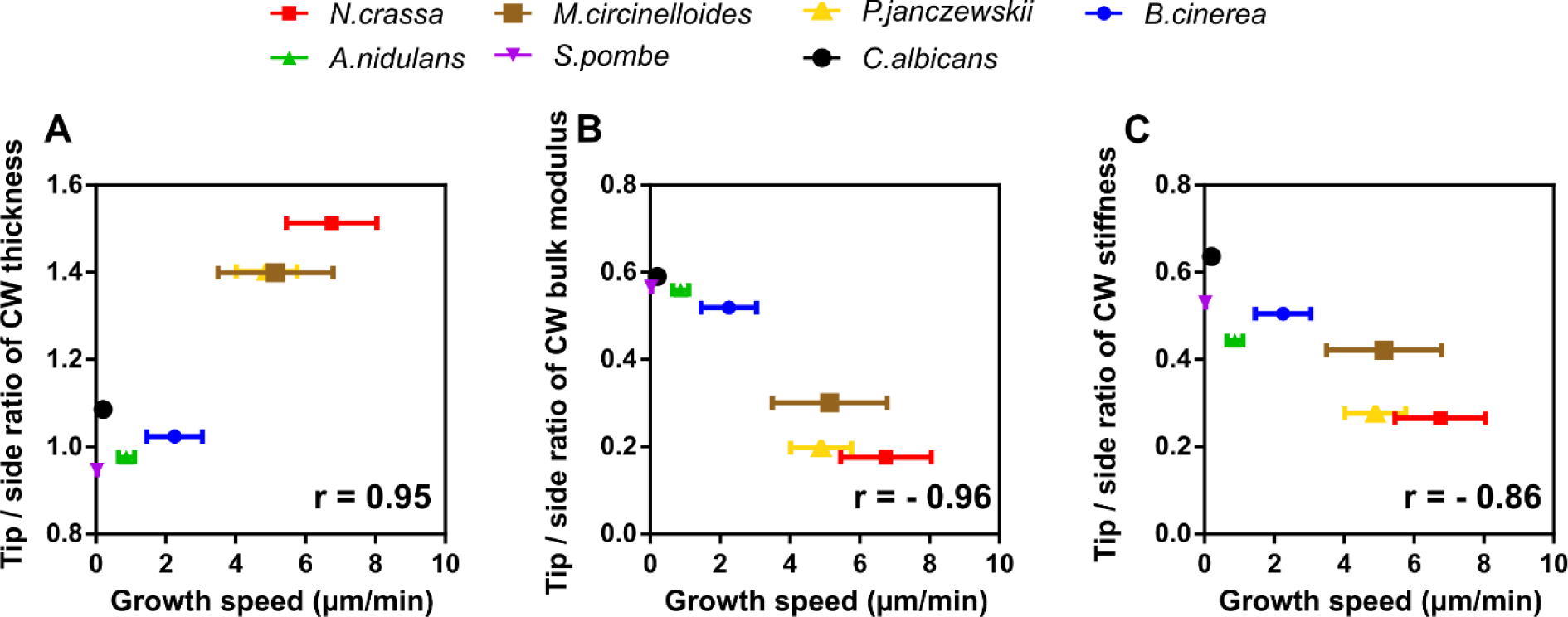
CW mechanical anisotropies and cell growth. Ratio of apical over lateral CW thickness (A), CW bulk modulus (B), and CW stiffness (C) plotted against mean growth speed. Error bars correspond to +/- S.D. Indicated r values on the graphs are Pearson correlation coefficients.

To understand if these properties may arise from constrains in tip CW tension, stress, or elastic strains, we computed these parameters at cell tips (Fig S4). We found that, CW tension at cell tips was negatively correlated with tip growth speeds, while CW stress did not show such a clear tendency (Fig S4A-B). The CW elastic strain was constant around ∼25% for most species and lower in some of the fastest cells (Fig S4C). The rescaled bending energy of tip CWs exhibited a large diversity of values, with a positive correlation with growth speeds (Fig S4D). Finally, previous modeling approaches have suggested that tip growth speed may depend on a product between an effective CW remodeling rate, *a*, and the elastic strain multiplied by the radius (PR^2^/hY) (24). We therefore used our mechanical measurements to extract the value of *a* as a function of tip growth speeds and found that faster growing cells may remodel their CWs much faster than slower ones, and that this parameter could be the most variable effector of tip growth speeds (Fig 3F). Together these results suggest that faster growing cells, exhibit less turgid cytoplasm, thin apical walls but higher CW remodeling rates; a strategy which may allow them to limit CW strains at cell tips and consequent risks of tip bursting.

In conclusion, we here implemented sub-resolution imaging and stress-strain measurements in a set of representative fungal species. These analyses allow to compute local thickness, bulk CW material properties, as well as turgor, and should be applicable to other types of fungi, allowing to assay cell mechanics in situations representative of infection phases, or response to antifungal agents, for instance. As such, these data provide an important resource of values for fungal cell mechanical parameters, that may impact our understanding of morphological variations found across species (34, 35). We found that CW bulk material properties vary across fungi, and are not necessarily scaled with cell size or growth speeds, which we interpret as a signature of variations in CW compositions or cross-links across fungi (32, 33). In contrast, CW thickness and turgor were inversely scaled with cell size and growth speeds, being both smaller in larger and more rapid growing hyphae. This inverse scaling tended to limit CW tension and elastic strains both on lateral and apical CWs. As shown in most fungi, hypo-osmotic treatments that effectively increase turgor, most often cause large tip-bursting and lysis phenotypes (23). Therefore, we propose that such mechanical scaling in filamentous fungi may arise from the selective pressure to limit CW elastic deformations, and promote CW integrity. Another important plausible consequence of lowering turgor, may also be at the level of membrane recycling. As shown in yeast, turgor opposes actin-based forces to set the rate of vesicle internalization (36–38). As very rapid tip-growing cells employ faster recycling rates, we speculate that reducing turgor could also be a mechanical strategy to facilitate trafficking events such as endocytosis in these cells (39). Future work coupling mechanical measurements with biochemical characterization of CWs and turgor regulation, may help addressing how biomechanical and biochemical designs have co-evolved to support morphogenetic diversity and survival.

## ACKNOWLEDGMENTS

We thank F. Leclerc (U. Paris Cité), R. Arkowitz (U. Nice), M. Peñalva (U. Madrid), A. Fleissner (U. Braunschweig), J. Dupont (MNHN) and P. Bassereau (I. Curie) for sharing strains, material and protocols, as well as A. Boudaoud for discussion. L.C. acknowledges the “École Doctorale LPI - Program Bettencourt” and a 4^th^ year fellowship from the Fondation de la Recherche Médicale (n°13171). This work was supported by the Centre National de la Recherche Scientifique (CNRS), the Université Paris Cité, and grants from La Ligue Contre le Cancer (EL2021.LNCC/ NiM), the European Research Council (ERC CoG “Forcaster” no. 647073), and the the Agence Nationale pour la Recherche (ANR, “CellWallSense” no. ANR-20-CE13-0003-02).

## AUTHOR CONTRIBUTIONS

Conceptualization, N.M., F.K. and L.C. Methodology, N.M., F.K., L.M. and L.C., Writing – Original Draft, N.M. and L.C. Draft Editing, N.M., and L.C.

## MATERIAL AND METHODS

### Fungal growth conditions and medium

Culture media were adapted to each species (see Table S2). Cells were grown on either Vogel’s medium (1.5% sucrose), MAE (2% malt and 2% glucose, 0.1% peptone), MCA (1% glucose and 5 mM ammonium tartrate), spider (1% nutrient broth and 1% mannitol), DO (2% glucose), or YE5S (3% glucose). Cultures were less than a day old, grown in liquid from spore solutions, using 8-well dishes (IBIDI GmbH, Martinriesd, Germany). *Candida albicans* was grown on liquid YPD (2% glucose), and cells in exponential growth were then harvested, rinsed, and placed in spider medium to induce filamentation. Experiments were done only on the hyphal form.

### Membrane and CW labeling

The CW was labeled using either 10µg/mL of WGA (Sigma, L9640) or 70µg/mL of ConA (ThermoFisher, C21421), depending on the species (see Table S2). For some non-tractable fungi, the plasma membrane was labeled using FM4-64 (ThermoFisher, T13320). To circumvent rapid endocytosis of the marker, droplets of ∼1µL of 1µg/mL of FM4-64 were added close to the chosen hyphae, yielding a bright membrane signal, and the images were taken within less than a minute after dye addition.

### Sorbitol treatments

Sorbitol shocks were done either in Ibidi wells or in PDMS chamber in which hyphal cells were maintained in the plane by being sandwiched in between dialysis membranes and coverglasses to constraint hyphae (40, 41). Cells were grown overnight from spore cultures in 1×4 cm PDMS chamber or in ibidi wells. Next, cells were imaged, rinsed with a medium supplemented with sorbitol, and imaged again less than a minute after. For *Mucor circinelloides* and *Penicillium janczewskii* a single layer of dialysis membrane was positioned between cells and the roof of the PDMS, in order to constrain hyphae to stay relatively flat, and limiting movements of hyphae after osmotic shocks.

For *N. crassa*, cells were grown on agar media for one day at 30°C in the dark, and a day at 4°C. An agar square with front line hyphae was then cut, and placed in an ibidi chamber with some liquid media (14). After the hyphae reached the border of the agar zone, images were taken, medium supplemented with sorbitol was added, and the same hyphae were imaged.

### Microscopy

Imaging was done at room temperature, using an inverted spinning-disk confocal microscope equipped with a motorized stage, automatic focus (Ti-Eclipse, Nikon, Japan), a Yokogawa CSUX1FW spinning unit, and a 100X oil-immersion objective (CFI Plan Apo DM 100X/1.4 NA, Nikon). We used either an EM-CCD camera (ImagEM-1K, Hamamatsu Photonics, Japan) with a 2.5X additional magnifying lens or a Prime BSI camera (Photometrics).

Laser ablation of the CW was done using a diffraction-limited spot from a 355 nm laser, with an iLas2 module (Gattaca, France) in the “Mosquito” mode, and performed on a 60X oil-immersion objective (CFI Apochromat 60X Oil lS, 1.4 NA, Nikon). Images were acquired with the 100X oil-immersion objective. The microscope was operated with the Metamorph software (Molecular Devices).

Tracking of cell shape during osmotic shocks was done using a Leica DMI6000 B microscope equipped with an A-Plan 40x/0.65 PH2 objective and an ORCA-Flash4.0LT Hamamatsu camera.

### Image analysis

#### CW thickness analyses

CW thickness measurements were done according to previous work (16, 24), using two-color mid-slice confocal images. Using an automated script, the positions of both signals were extracted by Gaussian fitting. Local distances between signals, which correspond to local values of cell wall thickness, were computed after chromatic shift and in/out signal correction.

#### Turgor pressure measurement

Turgor pressure was measured as in (16, 25), by comparing the shrinkage of cells rinsed with media supplemented with sorbitol at different concentrations (0-4M) with the shrinkage obtained by photo-ablation of the CW. Shapes were compared before and less than a minute after the osmotic shock to limit osmo-adaptation. Cell diameters were measured manually using Image J. The internal osmolyte concentration 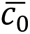 was measured as:

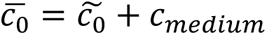

With 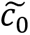 the sorbitol concentration at which the cell shrinks as much as with photoablation, and *c_medium_* the molarity of the medium used, measured using a vapor pressure osmometer (5600, Vapro® Vapor Pressure Osmometer).

From this, we calculated the effective concentration of the cytoplasm in the normal state 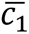:

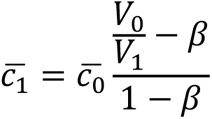

Where V_0_/V_1_ is the ratio between the volume before and after ablation, and *β* the inaccessible volume fraction, taken to be 0.22 (25).

This yields to an estimate of turgor, P, from the relationship:

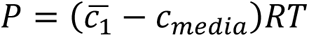

#### CW Young’s modulus

The Young’s modulus (Y) divided by pressure (P) was obtained by combining CW thickness measurement and photo-ablation assays. Assuming that the CW is homogenous and isotropic, the force balance in the cylindrical part yields:

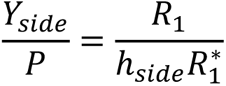

Where (R_1_) and (R_0_) are cell radii before and after photo-ablation respectively, and 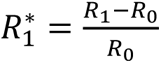, the elastic strain of the lateral CW.

For the hemispherical shape of tips, force balance leads to:

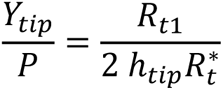

With 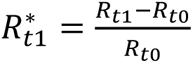 the elastic strain at cell tips, *R*_*t*1_ being the tip radius of curvature before ablation, and *R*_*t*0_ after. Importantly, these analytical formula were show previously to be valid for relatively long tubes through numerical simulations (21).

**Table S1.** Fungal species and strains used in this study. ^1^ André Fleissner, Technische Universität Braunschweig, Germany. ^2^ Florence Chapeland-Leclerc: UMR 8236, Université Paris cite, 75013 Paris France. ^3^ Joëlle Dupont, Muséum national d’Histoire naturelle, Paris France. ^4^ Miguel A Soto Peñalva: Centro de Investigaciones Biológicas Margarita Salas, Madrid 28040 Spain. ^5^ Robert Arkowitz : Institut de Biologie Valrose, Nice, France

| <u>Specie</u> | <u>Genotype</u> | <u>Source</u> |
| --- | --- | --- |
| *Neurospora crassa* | his-3::Ptef-GFP-Caax | Fleissner Andre <sup>1</sup> |
| *Mucor circinelloides* | WT | F. Leclerc Lab <sup>2</sup> |
| *Penicillium janczewskii* | WT | MNHN <sup>3</sup> |
| *Botrytis cinerea* | WT | F. Leclerc Lab |
| *Schizosaccharomyces pombe* | h+ ade6<<GFP-psy1 | Our lab<br>VD 55 |
| *Aspergillus nidulans* | pabaA1 yA2; argB2[argB*-gpdAp::GFP::PH-PLCdx2] | M Peñalva lab <sup>4</sup> |
| *Candida albicans* | pADHGFPctRac1 | R Arkowitz lab <sup>5</sup> |

**Table S2.** Fungal mechanical parameters, growth medium and labelling methods.

| Species | *N. crassa* | *B. cinerea* | *A. nidulans* | *P. janczewskii* | *C. albicans* (hyphae) | *S. pombe* | *M. circinelloides* |
| --- | --- | --- | --- | --- | --- | --- | --- |
| Medium | VMM | MAE | MCA | DO | spider | YE5S | DO |
| Radius (μm) | 4.6 | 2.51 | 1.61 | 3.1 | 1.19 | 1.98 | 3.93 |
| Growth speed (μm/h) | 405 | 135 | 52 | 293.4 | 12 | 1.74 | 308.4 |
| Tip CW Thickness (nm) | 72 | 118 | 76 | 94 | 123 | 170 | 83 |
| Lateral CW Thickness (nm) | 48 | 118 | 82 | 70 | 113 | 181 | 60 |
| Y/P tip | 87 | 33 | 61 | 82 | 16 | 23 | 140 |
| Y/P side | 494 | 66 | 120 | 413 | 30 | 40 | 443 |
| Sorbitol shock (M) to depressurize cells | 0.35 | 0.8 | 0.81 | 0.43 | >4 | 1.3 | 0.35 |
| Turgor Pressure (MPa) | 0.27 | 0.51 | 1.11 | 0.7 | 3 | 1.5 | 0.49 |
| Y tip (MPa) | 23 | 17 | 90 | 58 | 49 | 34.5 | 68 |
| Y side (MPa) | 133 | 33 | 134 | 289 | 90 | 60 | 217 |
| Membrane dye | GFP-CAAX | FM dye | GFP-PH | FM dye | Rac1-GFP | GFP-Psy1 | FM dye |
| CW dye | wga | conA | wga | wga | conA | GS-IB4 | wga |

### Supplemental figures and legends

**Figure S1:**
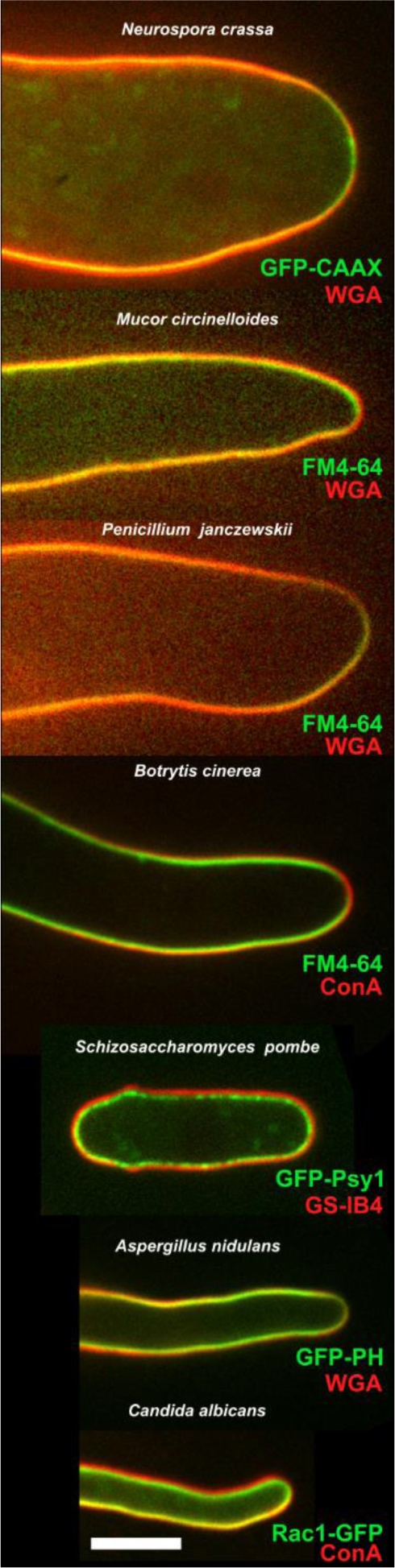
Labeling of the internal and external faces of the cell wall for cell wall thickness measurement. Examples of labeled cells, with in green the membrane labeling and in red the external CW labeling with lectins. WT strains membrane were labeled with FM-4 64 while tractable species were labeled with membrane associated domains or integral membrane proteins. The CW was labeled with either WGA, ConA or GS-IB4 lectins. Scale bar=5µm.

**Figure S2:**
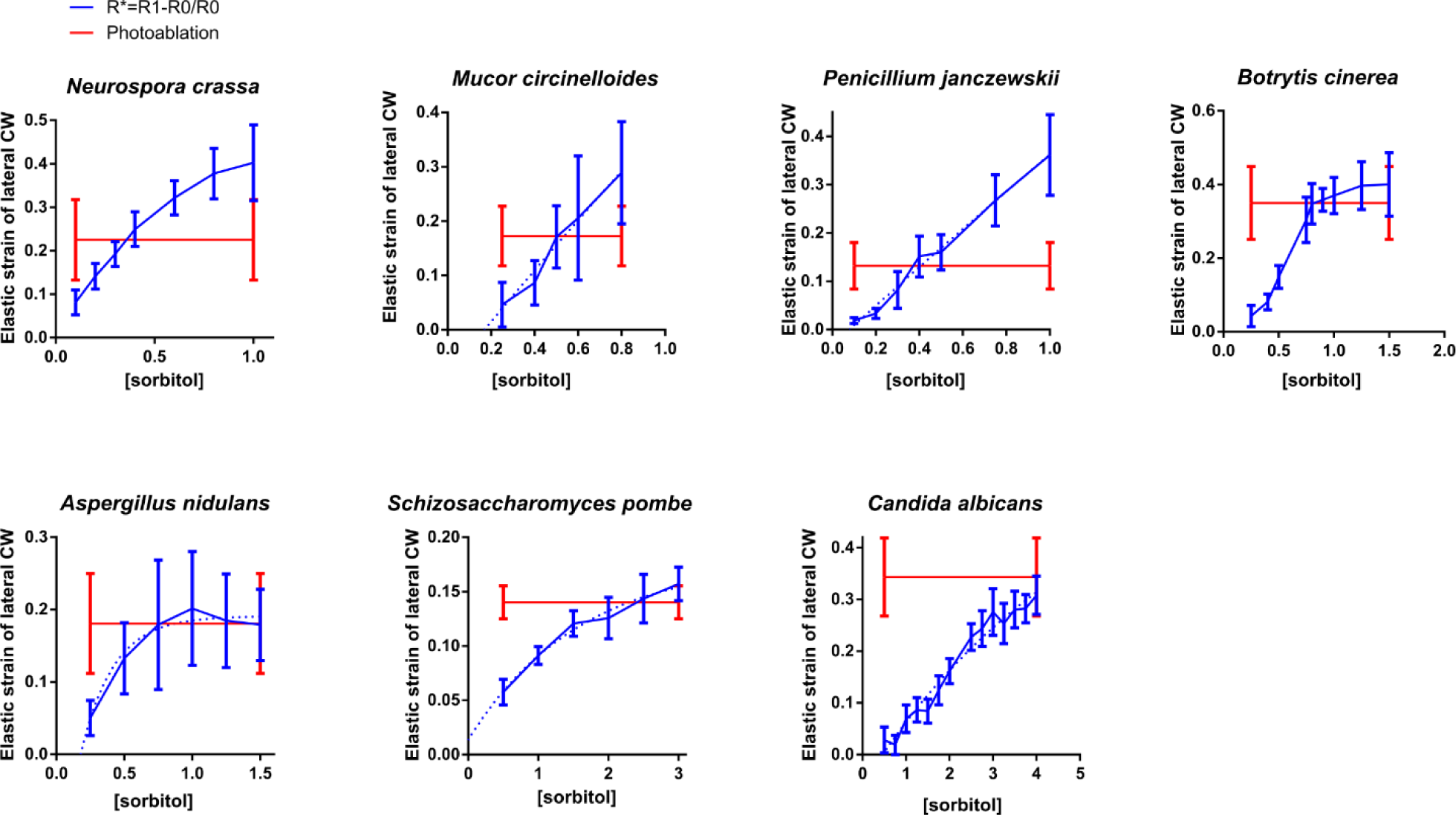
Turgor pressure measurement. Elastic strain of the lateral CW obtained for different sorbitol shocks compared to the one obtained by photo-ablation. The intersection of the two curves is used to estimate the external molarity needed to bring turgor to zero, and thus estimate turgor (n>15 for each species). Error bars correspond to +/- S.D.

**Figure S3:**
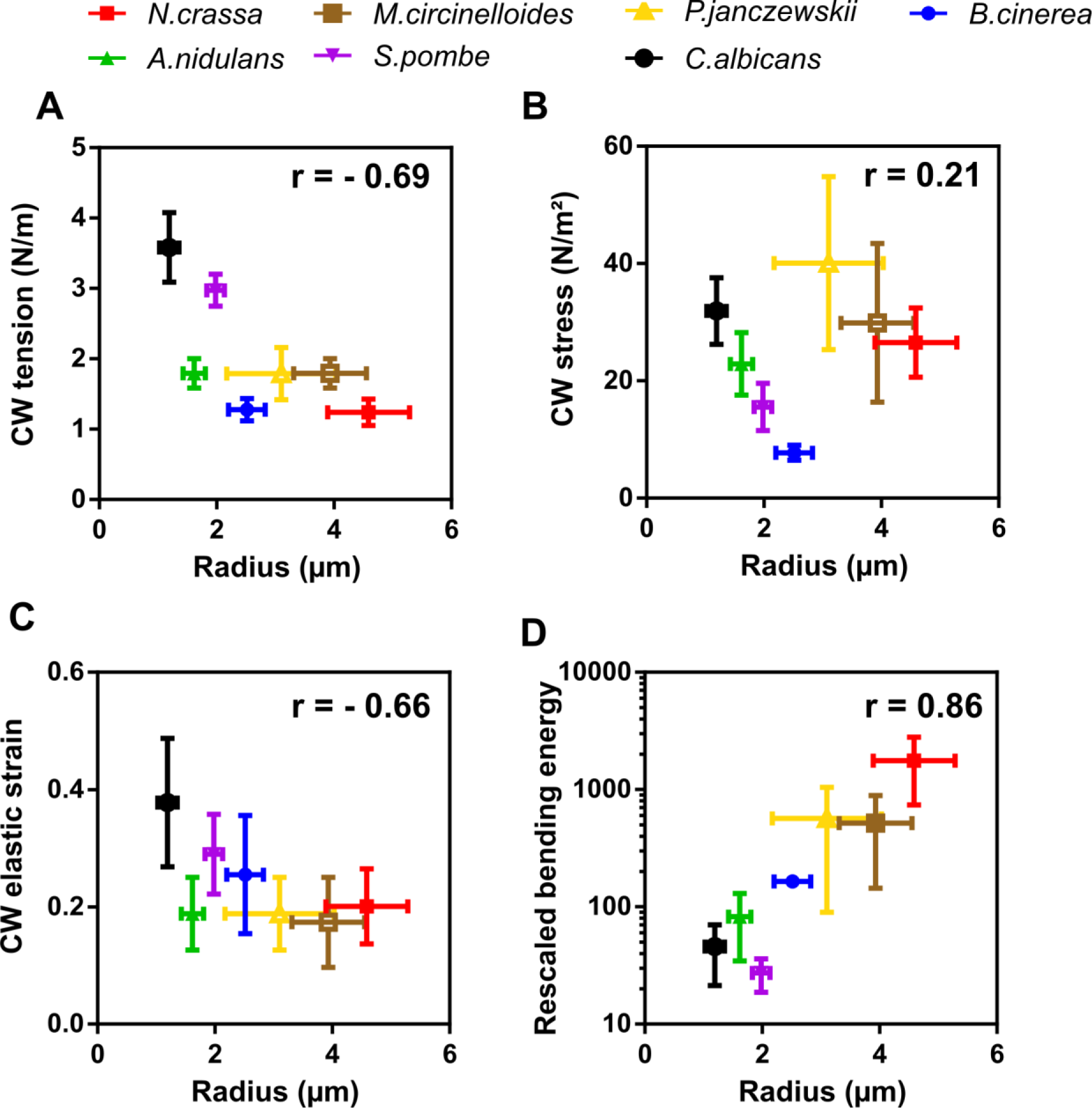
Scaling of lateral CW mechanical parameters with cell radius. (A) CW tension, (B) CW stress (C) CW elastic strain, and (D) rescaled bending energy versus cell radii (n>7 for each species). Error bars correspond to +/- S.D. Indicated r values on the graphs are Pearson correlation coefficients.

**Figure S4:**
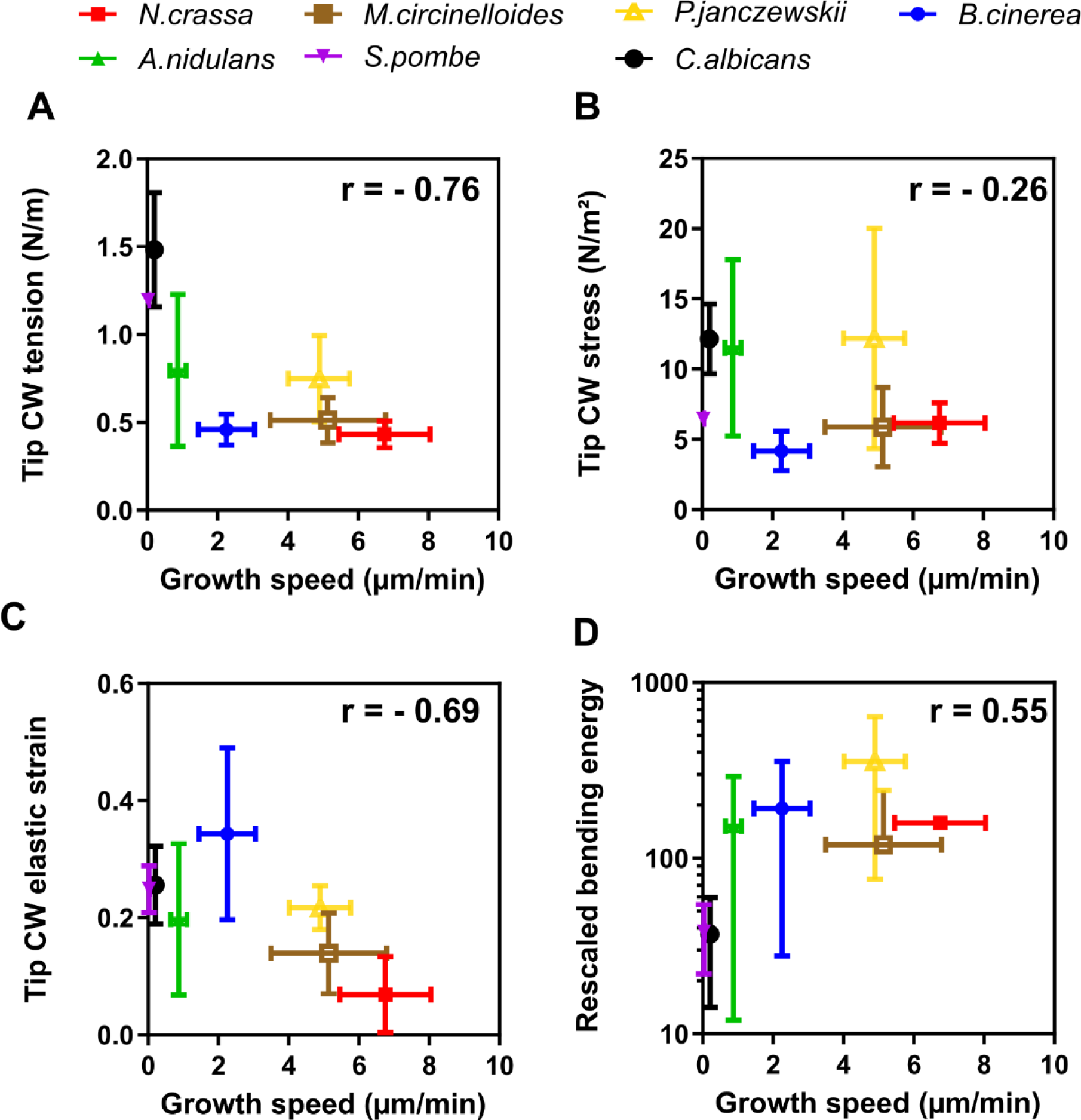
Scaling of tip CW mechanics with cell growth speeds. Tip CW. (A) tension, (B) stress, (C) elastic strain, and (D) rescaled bending energy versus cell growth speeds (n>9 for each species). Error bars correspond to +/- S.D. Indicated r values on the graphs are Pearson correlation coefficients.

## References

1. C. Municio-Diaz, et al., Mechanobiology of the cell wall – insights from tip-growing plant and fungal cells. Journal of Cell Science 135, jcs259208 (2022).

2. R. R. Lew, How does a hypha grow? The biophysics of pressurized growth in fungi. Nat Rev Microbiol 9, 509–18 (2011).

3. N. P. Money, Measurement of hyphal turgor. Experimental Mycology 14, 416–425 (1990).

4. R. R. Lew, N. N. Levina, S. K. Walker, A. Garrill, Turgor regulation in hyphal organisms. Fungal Genet Biol 41, 1007–15 (2004).

5. L. Beauzamy, N. Nakayama, A. Boudaoud, Flowers under pressure: ins and outs of turgor regulation in development. Annals of Botany 114, 1517–1533 (2014).

6. M. Bastmeyer, H. B. Deising, C. Bechinger, Force exertion in fungal infection. Annu Rev Biophys Biomol Struct 31, 321–41 (2002).

7. R. Mishra, N. Minc, M. Peter, Cells under pressure: how yeast cells respond to mechanical forces. Trends in Microbiology (2022). 10.1016/j.tim.2021.11.006.

8. M. A. Naranjo-Ortiz, T. Gabaldón, Fungal evolution: cellular, genomic and metabolic complexity. Biol Rev Camb Philos Soc 95, 1198–1232 (2020).

9. G. Steinberg, M. A. Penalva, M. Riquelme, H. A. Wosten, S. D. Harris, Cell Biology of Hyphal Growth. Microbiol Spectr 5 (2017).

10. V. Davi, N. Minc, Mechanics and morphogenesis of fission yeast cells. Curr Opin Microbiol 28, 36–45 (2015).

11. S. G. Martin, R. A. Arkowitz, Cell polarization in budding and fission yeasts. FEMS Microbiol Rev 38, 228–53 (2014).

12. M. C. Ruiz-Roldán, et al., Nuclear Dynamics during Germination, Conidiation, and Hyphal Fusion of Fusarium oxysporum. Eukaryot Cell 9, 1216–1224 (2010).

13. S. Fischer-Parton, et al., Confocal microscopy of FM4-64 as a tool for analysing endocytosis and vesicle trafficking in living fungal hyphae. Journal of Microscopy 198, 246–259 (2000).

14. R. López-Franco, S. Bartnicki-Garcia, C. E. Bracker, Pulsed growth of fungal hyphal tips. Proceedings of the National Academy of Sciences 91, 12228–12232 (1994).

15. S. Taheraly, D. Ershov, S. Dmitrieff, N. Minc, An image analysis method to survey the dynamics of polar protein abundance in the regulation of tip growth. J Cell Sci 133, jcs252064 (2020).

16. L. Chevalier, et al., Cell wall dynamics stabilize tip growth in a filamentous fungus. PLOS Biology 21, e3001981 (2023).

17. M. Köhli, V. Galati, K. Boudier, R. W. Roberson, P. Philippsen, Growth-speed-correlated localization of exocyst and polarisome components in growth zones of Ashbya gossypii hyphal tips. Journal of Cell Science 121, 3878–3889 (2008).

18. F. D. Kelly, P. Nurse, Spatial control of Cdc42 activation determines cell width in fission yeast. Mol Biol Cell 22, 3801–11 (2011).

19. D. Bonazzi, et al., Actin-Based Transport Adapts Polarity Domain Size to Local Cellular Curvature. Curr Biol 25, 2677–83 (2015).

20. N. Minc, A. Boudaoud, F. Chang, Mechanical forces of fission yeast growth. Curr Biol 19, 1096–101 (2009).

21. V. Davi, et al., Systematic mapping of cell wall mechanics in the regulation of cell morphogenesis. Proc Natl Acad Sci U S A 116, 13833–13838 (2019).

22. J. D. Stenson, P. Hartley, C. Wang, C. R. Thomas, Determining the mechanical properties of yeast cell walls. Biotechnol Prog 27, 505–12 (2011).

23. S. Bartnicki-Garcia, E. 1972 Lippman, The Bursting Tendency of Hyphal Tips of Fungi: Presumptive Evidence for a Delicate Balance between Wall Synthesis and Wall Lysis in Apical Growth. Microbiology 73, 487–500 (1972).

24. V. Davì, et al., Mechanosensation Dynamically Coordinates Polar Growth and Cell Wall Assembly to Promote Cell Survival. Developmental Cell 45, 170–182.e7 (2018).

25. E. Atilgan, V. Magidson, A. Khodjakov, F. Chang, Morphogenesis of the Fission Yeast Cell through Cell Wall Expansion. Curr Biol 25, 2150–7 (2015).

26. R. Vauchelles, D. Stalder, T. Botton, R. A. Arkowitz, M. Bassilana, Rac1 Dynamics in the Human Opportunistic Fungal Pathogen Candida albicans. PLOS ONE 5, e15400 (2010).

27. A. Serrano, et al., Spatio-temporal MAPK dynamics mediate cell behavior coordination during fungal somatic cell fusion. J Cell Sci 131, jcs213462 (2018).

28. K. Youssef, S. R. Roberto, A. G. de Oliveira, Ultra-Structural Alterations in Botrytis cinerea—The Causal Agent of Gray Mold—Treated with Salt Solutions. Biomolecules 9, 582 (2019).

29. A. P. Trinci, A. J. Collinge, Hyphal wall growth in Neurospora crassa and Geotrichum candidum. J Gen Microbiol 91, 355–361 (1975).

30. L. Zhao, et al., Elastic Properties of the Cell Wall of Aspergillus nidulans Studied with Atomic Force Microscopy. Biotechnology Progress 21, 292–299 (2005).

31. S. Bartnicki-Garcia, W. J. Nickerson, Isolation, composition, and structure of cell walls of filamentous and yeast-like forms of Mucor rouxii. Biochim Biophys Acta 58, 102–119 (1962).

32. N. A. R. Gow, J.-P. Latge, C. A. Munro, The Fungal Cell Wall: Structure, Biosynthesis, and Function. Microbiol Spectr 5 (2017).

33. S. J. Free, Fungal cell wall organization and biosynthesis. Adv Genet 81, 33–82 (2013).

34. M. E. Ohairwe, B. D. Živanović, E. R. Rojas, A fitness landscape instability governs the morphological diversity of tip-growing cells. Cell Reports 113961 (2024). 10.1016/j.celrep.2024.113961.

35. O. Campàs, E. Rojas, J. Dumais, L. Mahadevan, Strategies for cell shape control in tip-growing cells. American Journal of Botany 99, 1577–1582 (2012).

36. R. Basu, E. L. Munteanu, F. Chang, Role of turgor pressure in endocytosis in fission yeast. Mol Biol Cell 25, 679–687 (2014).

37. S. Dmitrieff, F. Nédélec, Membrane Mechanics of Endocytosis in Cells with Turgor. PLoS Comput Biol 11, e1004538 (2015).

38. R. Ma, J. Berro, Endocytosis against high turgor pressure is made easier by partial coating and freely rotating base. Biophysical Journal 120, 1625–1640 (2021).

39. A. Picco, C. P. Toret, A.-S. Rivier-Cordey, M. Kaksonen, An evolutionary cell biology perspective into the diverging mechanisms of clathrin-mediated endocytosis in dikarya fungi. [Preprint] (2024). Available at: https://www.biorxiv.org/content/10.1101/2024.03.28.587219v1 [Accessed 16 April 2024].

40. G. Charvin, F. R. Cross, E. D. Siggia, A microfluidic device for temporally controlled gene expression and long-term fluorescent imaging in unperturbed dividing yeast cells. PLoS One 3, e1468 (2008).

41. A. Haupt, D. Ershov, N. Minc, A Positive Feedback between Growth and Polarity Provides Directional Persistency and Flexibility to the Process of Tip Growth. Curr Biol 28, 3342–3351 e3 (2018).

42. T. Y. James, et al., Reconstructing the early evolution of Fungi using a six-gene phylogeny. Nature 443, 818–822 (2006).

43. J. E. Stajich, et al., The Fungi. Current Biology 19, R840–R845 (2009).

